# Reward sensitivity following boredom and cognitive effort: A high-powered neurophysiological investigation

**DOI:** 10.1101/177220

**Authors:** Marina Milyavskaya, Michael Inzlicht, Travis Johnson, Michael Larson

## Abstract

What do people feel like doing after they have exerted cognitive effort or are bored? Here, we empirically test whether people are drawn to rewards (at the neural level) following cognitive effort and when bored. This elucidates the experiences and consequences of engaging in cognitive effort, and compares it to the consequences of experiencing boredom, an affective state with predicted similar motivational consequences. Event-related potentials were recorded after participants (N=243) were randomized into one of three conditions – boredom (observing strings of numbers), cognitive effort (adding 3 to each digit of a four-digit number), or control. In the subsequent task, we focused on the feedback negativity (FN) to assess the brain’s immediate response to the presence or absence of reward. Phenomenologically, participants in the boredom condition reported more fatigue than those in the cognitive effort condition. Results suggest participants in the boredom condition exhibited larger FN amplitude than participants in the control condition, while the cognitive effort condition was neither different from boredom nor control. The neural and methodological implications for ego depletion research, including issues of replicability, are discussed.

## Reward sensitivity following boredom and cognitive effort: A high-powered neurophysiological investigation

Imagine you just spent the morning grading exams for a large course. It was boring work, and you feel drained. What do you feel like doing? Relaxation might immediately come to mind, but what if you have to get back to work? You may find other ways of rewarding yourself – perhaps eating a chocolate bar, allowing yourself a few minutes to peruse Facebook, or generally engaging in some other activity that you find pleasurable or rewarding. Although people generally seek out rewarding experiences, here we wonder if this is especially the case after being bored or engaging in cognitive effort. In the present study, we empirically test whether people are more drawn to rewards after boredom or after engaging in cognitive effort compared to when they are neither, by examining the sensitivity to rewards on a neural level.

### Cognitive effort, ego depletion, and shifting priorities towards rewards

Neurocognitive accounts of cognitive effort view it as the mobilization of resources necessary to attain a desired level of performance (Shenhav et al., 2017). The term cognitive effort is frequently used interchangeably with cognitive control, with some suggesting that cognitive effort drives the decision to engage control (Westbrook & Braver, 2015). Cognitive effort, and in particular decisions about engaging effort, is thought to be neurally mediated by the dorsal anterior cingulate cortex (dACC) and the lateral prefrontal cortex (lPFC; see Shenhav et al., 2017, for review). Importantly, cognitive effort is generally considered inherently aversive or costly (Kool et al. 2010; Westbrook, Kester & Braver, 2013). According to some recent models, this cost is then weighed against the benefits of exerting the effort. Although there are some differences in the accounts of the reasons why mental effort is costly (see Shenhav et al., 2017, for a review), one proposed suggestion relates to the notion of opportunity costs (Kurzban et al., 2013). When engaging in cognitive effort on any given task, a person foregoes opportunities to engage in other (potentially valuable) tasks – these lost opportunities are termed the opportunity cost of persisting at an effortful task. According to this model, the costs of engaging in further effort are expected to rise further as more effort is exerted, with the value of effort exertion (i.e., the cost-benefit ratio) diminishing proportionally (Kurzban et al., 2013). After incurring large costs, people may want a (proportionally large) reward.

The consequences of exerting cognitive effort have also been the focus of a large body of literature on ego depletion. Ego depletion refers to a psychological state whereby people feel unable or unwilling to exert effort following an effortful task. It is akin to a state of mental fatigue (Inzlicht, Schmeichel, & Macrae, 2014), whereby after engaging in an activity that requires effortful control, people perform more poorly on a second task, also requiring effortful control (Baumeister, Bratslavsky, Muraven, & Tice, 1998). This sequential ego depletion paradigm (consisting of two sequential tasks requiring cognitive effort) has been used in hundreds of studies (see Hagger et al. 2010), although a recent pre-registered replication report did not find an effect for one specific operationalization of depletion (Hagger et al. 2016).

While the very existence and magnitude of the ego depletion effect are currently being examined (Carter & McCullough, 2014; Hagger et al. 2016), some have wondered how replication difficulties are affected by the possible mechanisms underlying ego depletion (Inzlicht et al. 2014; Kurzban, Duckworth, Kable, & Myers, 2013). One explanation for the “depletion” period so frequently observed after a demanding self-control task centers on motivation – after exerting effort on one task, people are no longer motivated to exert further effort on a subsequent task, and instead prefer to ‘indulge’ in an immediate temptation (e.g., of slacking off, venting one’s anger, eating the delicious food, etc.). According to this shifting priorities model of self-control (Inzlicht et al. 2014; Milyavskaya & Inzlicht, 2017), this shift in motivation would be accompanied by shifts in attention, perception, emotion, and memory. Thus after an effortful task, people may be more drawn to rewards, and may more readily notice opportunities to indulge in rewarding behaviours. Fatigue is thought to have similar motivational consequences (Hockey, 2013).

For example, participants are more likely to gamble (Bruyneel, Dewitte, Franses, & Dekimpe, 2009), shop (Vohs & Faber, 2007), eat (Vohs & Heatherton, 2000), and smoke (Shmueli & Prochaska, 2009) after engaging in cognitive effort. Similarly, an experience-sampling study found that people were more likely to succumb to tempting desires when depleted (Hofmann, Vohs & Baumeister, 2012). However, since enactment of a temptation is jointly determined both by the strength of the desire and the amount of effort exerted (Hofmann et al., 2012), such effects could occur either because participants are more sensitive/tempted by the rewards (i.e., the desire is greater; Schmeichel, Harmon-Jones, & Harmon-Jones, 2010), or because they exert less self-control when faced with the desire.

To our knowledge, only a few studies have examined the effects of ego depletion or the exertion of cognitive effort on the perception of rewards. In one study, participants who were depleted were more accurate in detecting a reward-related symbol ($) than non-reward symbol (&) in rapidly presented images (Schmeichel et al., 2010). In another particularly relevant study, dieters who were depleted exhibited greater food-cue-related activity in areas of the brain associated with coding reward values, and other areas associated with self-control (Wagner, Altman, Boswell, Kelley & Heatherton, 2013). In that study, chronic dieters either completed a depletion task or a control task and then viewed desirable foods while in an fMRI scanner. Participants in the depletion condition had greater activation in the orbitofrontal cortex, as well as greater functional connectivity between the orbitofrontal cortex and the inferior frontal gyrus, than those in the control condition (Wagner et al., 2013). Together, these studies support the possibility that after people exert cognitive effort, they exhibit increased sensitivity towards rewards.

### Boredom as similar motivational state

If the consequences of effort expenditure affect states of motivation, then depletion might have similar motivational properties as other states that influence motivation towards rewards. One such state is boredom. Boredom is typically described as an affective state that results from the inability to “successfully engage attention with internal or external information” (Eastwood, Frischen, Fenske, & Smilek, 2012, pg. 484); it is characterized by “core motivational deficits accompanied by a phenomenological experience of a lack of interest or affective engagement.” (Goldberg, Eastwood, Laguardia & Danckert, 2011, pg. 649).

Although it may at first seem paradoxical that engaging in cognitive effort would lead to the same consequences as boredom, there are several reasons to theorize that boredom might have similar motivational consequences to depletion. First, research on vigilance tasks that require participants to monitor displays for infrequent stimuli for a prolonged period of time (e.g., Mackworth, 1948) have been alternatively interpreted as inducing fatigue (i.e., depletion) or boredom (Pattyn, Neyt, Henderickx, & Soetens, 2008). Indeed, research has shown that both the “depletion of information-processing resources” (i.e., an overload of the attentional system; Pattyn et al. 2008, pg. 377) and under-arousal both lead to a decrease in vigilance (see Pattyn et al. 2008). In other words, vigilance tasks might induce both (1) fatigue and (2) boredom, both of which might have similar downstream consequences on subsequent behavior. Second, animal models of boredom find that animals housed in cages with no opportunities for enrichment display more interest in novel stimuli and consume more food rewards (Meagher & Mason, 2012) – that is, these ‘bored’ animals are more attuned to rewards. Similarly, in humans greater and more frequent experiences of boredom have been linked to engaging in more impulsive, reward-seeking behaviour including gambling (Blaszczynski et al. 1990), overeating and binge eating (Abramson & Stinson, 1977; Myhre et al. 2015), and alcohol and drug abuse (Iso-Ahola & Crowley, 1991); this is similar to the impulsive tendencies of depleted participants (e.g., Vohs & Heatherton, 2000; Vohs & Faber, 2007). While research finds that people who are generally prone to boredom engage more frequently in such impulsive behaviours, to our knowledge there has not been any research examining whether state boredom would directly affect a person’s orientation towards rewards.

One proposed function of boredom is that boredom serves as an indicator to pursue an alternate goal (Bench & Lench, 2013). A similar motivational function of depletion has also been proposed, with depletion or fatigue seen as a stop-signal, a signal to end cognitive effort and engage in other pursuits (Hockey, 2013; Inzlicht et al., 2014; Kurzban et al. 2013). This suggests that boredom and depletion might have similar functions in orienting humans to disengage from current behaviour and seek other (more rewarding) alternatives. A purpose of the current study, then, is to test whether boredom also elicits a stronger orientation towards rewards. Specifically, we predicted that participants who were either bored or depleted would show an increased sensitivity to rewards compared to participants who were neither depleted nor bored.

### The Feedback Related Negativity (FN)

In the present study, we looked at reward sensitivity as the brain’s immediate responses to reward using electroencephalographic (EEG) recordings, focusing on the feedback negativity (FN) component of the scalp-recorded event-related potential (ERP). The FN is a negative deflection in the ERP at frontocentral electrode recording sites that occurs around 250ms after feedback presentation and is larger (i.e., more negative) to unfavorable than favorable outcomes. Recent research (see Proudfit, 2015 for review) suggests that the FN may represent a positive response to positive feedback (reflecting gains or rewards) that is reduced following negative feedback, with the difference between negative and positive feedback (negative minus positive) representing the negative deflection seen in the FN. In this view, the FN is actually a decrease in the positivity associated with reward/favorable feedback (Proudfit, 2015), rather than a negative deflection per se. Nonetheless, The FN has good psychometric properties (Levinson et al., 2017) and has been correlated with subjective interest in rewards (Bress & Hajcak, 2013), and with approach motivation more generally (Threadgill & Gable, 2016). In the present study, we examine the FN to evaluate the extent to which depleted and bored participants are drawn towards rewards.

### Present study

We conducted a high-powered study to examine the effects of effort expenditure and boredom on reward sensitivity. We also included measures of phenomenology as a manipulation check, expecting that the effort condition would result in high self-reported effort and fatigue (compared to the boredom condition), and that the boredom condition would lead to greater boredom (compared to the effort condition). For our main research questions, we first hypothesized that participants who exert cognitive effort (i.e., the effort condition) would have a greater sensitivity to rewards (as indexed by the FN) than participants in the control condition.

Importantly, we expected the boredom condition to have an effect similar to the effort condition, with both resulting in higher sensitivity to rewards than the control condition. We also conducted exploratory analyses to examine whether the type of reward mattered. Specifically, previous research has found that monetary rewards elicited greater FN than no-value rewards; while we expected to replicate this main effect, we did not have specific predictions as to whether there would be an interaction with condition.

## Method

### Participants

There is no consensus on the typical effect size for the sequential task paradigm (although the effect is likely small, if it exists at all; see Inzlicht & Berkman, 2015); more importantly, since we used a neural index of sensitivity to rewards rather than a measure of control, we had nothing to guide our selection of expected effect size. We thus aimed for a medium effect. A power analysis indicated that 210 participants would allow the detection of a medium effect (f =.25) in a one-way ANOVA with 3 groups with a power of .90 (G*Power). We set out to recruit at least 210 participants, but continued recruiting until the end of the academic semester. Participants were 243 university students who completed the study for course credit - a sample that is an order of magnitude larger than typical EEG studies. Thirty-one participants were excluded from the experiment due to equipment malfunction or too much noise in the data, resulting in unusable EEG data (21 participants), or insufficient artifact-free trials (10 participants; see explanation below). The final sample consisted of 212 participants (41.9%female) ages 18-26 (M = 20.84, SD = 2.09). All participants were healthy, free from neurological disease, and had no food allergies.

### Procedure

Participants came into the lab and were connected to the EEG system. They were then randomly assigned to one of three conditions, completing either one of two numbers tasks (effort and boredom conditions) followed by the computerized door task (designed to elicit the FN), or, in the neutral condition, went straight to the computerized doors task. For participants in the effort and boredom conditions, the computerized door task was presented as a separate study: “Since you are already connected to the EEG, and our study is not very long, we have partnered with another researcher to help them pilot a new task, so we ask that you do this task now.” After the door task, participants completed personality questionnaires (included in the study for exploratory purposes; see https://osf.io/e39as/?view_only=03a768b3ec084a86a4a1ce488ce6d164 for all materials). Participants in the control condition completed the same numbers task as those in the effort condition *after* all other study procedures were completed (to equate participation length, as required by the institution’s IRB board). At the end of the study, participants were given their additional monetary and candy reward and debriefed. The outline of the procedure is illustrated in Figure 1.

**Figure 1.**
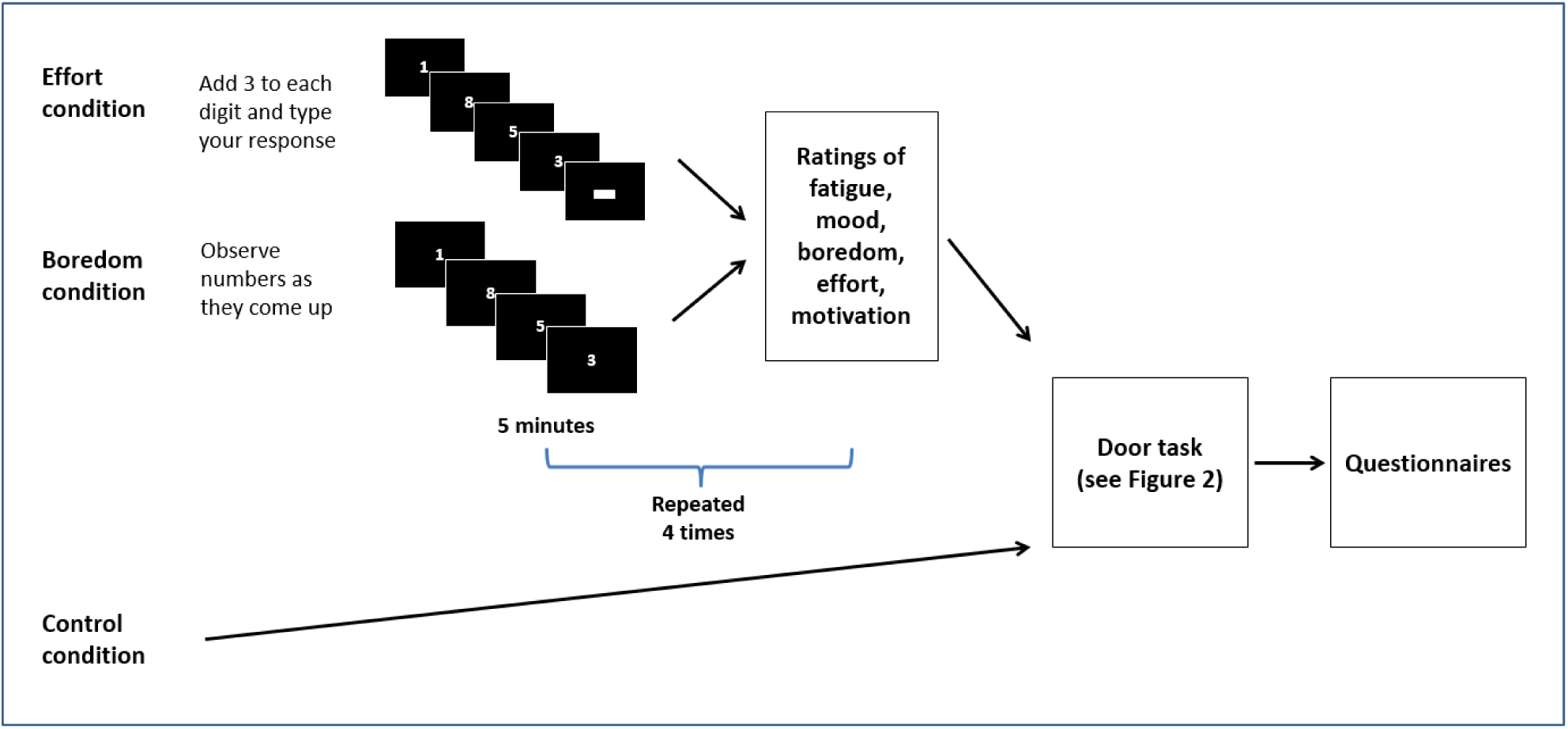
The procession of experimental tasks across the three conditions. Participants in the effort and boredom conditions completed 20 minutes of either the add 3 task or the passive number viewing task in four blocks of five minutes each; each block was followed by self-reported ratings. Participants in all three conditions then proceeded to the door task followed by questionnaires.

### Experimental manipulation

Participants were randomly assigned using a random number generator into one of three conditions. In the effort and boredom conditions, participants completed a 20-minute task involving either number manipulation or passive number viewing. In both tasks, participants were presented with strings of four digits one at a time. In the effort condition, participants were asked to add 3 to each digit and type their response (e.g., 9234 results in 2567; Kahneman, 2011). This is a task that requires high cognitive effort, result in maximal pupil dilation, and is experienced as subjectively difficult (Kahneman, 2011). In the boredom condition, participants were asked to simply observe the numbers as they came up. Since no active effort is required other than simple vigilance, we expected that participants would experience a lack of engagement and stimulation, resulting in under-arousal characteristic of boredom. Both tasks consisted of 4 blocks of 5 minutes each, for a total of 20 minutes. After each block, participants were asked to rate their fatigue, mood, boredom, effort, and motivation using a visual-analog scale (VAS) for each item: “How are you feeling right now” (Fatigued-Energized); “What is your mood right now” (Unpleasant-Pleasant); “How are you feeling right now?” (Bored-Interested); “How hard are you trying to do well on the task” (Not trying very hard – trying very hard); “How much do you feel like you are participating in this study” (Because I feel like I have to – Because it is personally important to me). The VAS scales were translated into a number from 0 to 100, with larger numbers representing the labels on the right.

*Computerized door task*

All participants then completed the door lottery task. This task was modeled on a similar task commonly used to elicit the FN (Proudfit, 2015; Weinberg, Riesel & Proudfit, 2014), but with one important difference: we added a third reward option consisting of a food reward (M&M candy). Since food, and in particular sugar/glucose, has been conceptualized by both lay people and some researchers as an effective antidote to depletion (Baumeister & Tierney, 2011; Gailliot et al., 2007; cf. Kurzban, 2010), we were interested in whether these lay theories would translate into people being more oriented towards such food-related rewards, such that M&M candy would be especially rewarding to someone who has recently exerted effort.

Our task consisted of 144 trials organized in 4 blocks to provide participants with breaks (see Figure 2). On each trial, participants saw two doors and were asked to choose one to open. Once the door was opened, participants received feedback to indicate whether they won (green arrow pointing up) or lost (red arrow pointing down). Prior to each trial, participants were informed whether on the upcoming trial they would have the chance to earn or lose money, candy (M&Ms), or nothing. There were 48 trials for each type of reward (randomized within-block), with positive feedback presented on 50% of the trials. Prior to the task, participants were informed that they would receive 50c for each ‘correct’ money trial and 2 M&M candies for each ‘correct’ candy trial, and lose 25c on each ‘incorrect’ money trial or 1 M&M candy on each incorrect’ candy trial (due to losses looming larger than gains; see Proudfit, 2015). The order and timing of all stimuli are as follows: (i) one of three texts (‘Click for the next round’, ‘Click for a chance to win money’ or ‘Click for a chance to win candy’) appeared until the response is made; (ii) the graphic of two doors is presented until a response is made, (iii) a fixation mark is presented for 1000 ms, (iii) a feedback arrow is presented for 2000 ms, (iv) a fixation mark is presented for 1500 ms. In between each block, participants were told how much money and M&Ms they earned in the previous block (this was constant for all participants).

**Figure 2.**
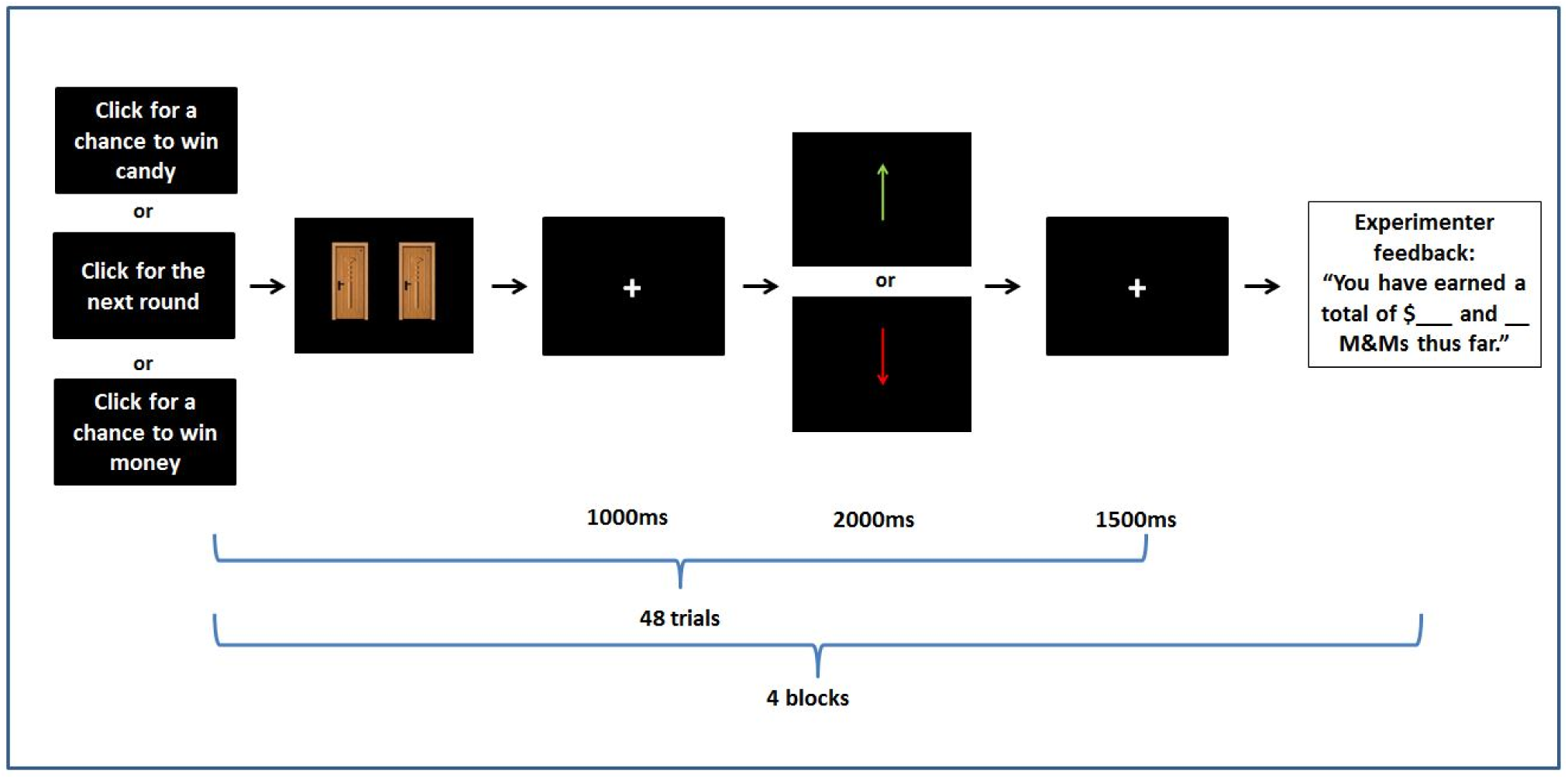
The computerized door task. Participants completed four blocks of the door task, each consisting of 36 trials. Before each trial participants were informed whether on the upcoming trial they would have the chance to earn or lose money, candy (M&Ms), or nothing. Participants then saw two doors and were asked to choose one to open. Once the door was opened, participants received feedback to indicate whether they won (upwards green arrow) or lost (downward red arrow); positive feedback was presented on 50% of the trials. Participants received 50c or 2 M&M candies for each ‘correct’ trial, and lost 25c or 1 M&M candy on each ‘incorrect’ trial. EEG signals were time-locked to the presentation of the feedback (upwards green arrow or downward red arrow).

### EEG Data Acquisition and Reduction

Electroencephalogram data was recorded during the doors task from 128 scalp sites using a geodesic sensor net and Electrical Geodesics, Inc., (EGI; Eugene, Oregon) amplifier system (20K gain, nominal bandpass=.10-100Hz). Electrode placements enabled recording vertical and horizontal eye movements reflecting electro-oculographic (EOG) activity. Data from the EEG was referenced to Cz and digitized continuously at 250Hz with a 16-bit analog-to-digital converter. A right posterior electrode approximately two inches behind the right mastoid served as common ground. Electrode impedance was maintained below 50k?. EEG signals were time-locked to the presentation of the feedback (upwards green arrow or downward red arrow). EEG data was segmented off-line into epochs between -200 ms before and 800 ms after stimulus presentation with a 200ms baseline correction. Data were filtered with 0.1 to 30Hz bandpass. We removed eye blinks and saccades using independent components analysis (ICA) implemented in the ERP PCA Toolkit (Dien, 2010). The ICA components that correlated at .9 with the scalp topography of two templates, one generated based on the current data and another provided by the ERP PCA Toolkit author, were removed. Trials were considered bad if more than 15% of channels were marked bad. Channels were marked bad if the fast average amplitude exceeded 100µV or if the differential average amplitude exceeded 50µV.

Previous work with high-density arrays suggests averaging across electrodes provides increased reliability (Baldwin et al. 2015). Thus, from the artifact-free epochs we extracted the ERPs within the time window from 175ms to 225ms across an average of 7 frontal electrodes including electrodes 5, 6 (FCz), 7, 12, 13, 106, and 112 (see Larson et al. 2010 for figure with electrode locations). This time window was selected using the *collapsed localizers* approach suggested by Luck and Gaspelin (2016) to deal with the problem of multiple implicit comparisons inherent in ERP research.

In the collapsed localizer approach for this study, we averaged all the waveforms across all conditions and selected the time window that demonstrated the largest negative deflection at the general time frame of interest. This ensured that we did not select a window based on visual differences between conditions (see Luck & Gaspelin, 2016). To obtain the general FN, the average of all ‘correct’ feedback trials was subtracted from the average of all incorrect feedback trials (irrespective of reward type). More negative numbers indicate a stronger FN. Then, to obtain the FN for each specific reward type (for use in multilevel analyses to examine differences by type of reward) the average of all ‘correct’ feedback trials for each given reward type was subtracted from the average of all incorrect feedback trials for that reward type. For these, only instances with 10 or more artifact-free trials of that type were used, such that some participants did not have data for one or two of the reward types.

ERP amplitude estimates were determined to be reliable with a minimum of 10 trials per condition (see Clayson et al. 2013). As reliability is dependent on each specific sample and study), dependability estimates (a generalizability theory [G-theory] analogue of reliability) were calculated for each group and condition. Using formulas provided by Baldwin et al. (2015) in the ERP Reliability Analysis (ERA) Toolbox v 0.3.2 (Clayson & Miller, in press). The ERA Toolbox calculated ERP dependability based on algorithms from generalizability theory (see Baldwin et al., 2015 for review) and used CmdStan v 2.10.0 to implement the analyses in Stan. When each condition had at least 10 trials, point dependability estimates all exceeded .70 except 1 (Effort Neutral Correct trials = .67). Dependability estimates, 95% credible intervals, mean numbers of trials, and trial count range as a function of group and condition are presented in Table 1. These findings do not contradict those of Levinson et al., (2017) who suggest a minimum of 20 trials is needed for a reliable FN because reliability/dependability are specific to each individual sample and this particular sample simply required fewer trials to reach dependability than the Levinson study.

**Table 1.**

|  | Dependability | 95% CI | Mean # Trials | Trial Range |
| --- | --- | --- | --- | --- |
| Boredom Candy Correct | .85 | .80; .90 | 20.2 | 12; 24 |
| Boredom Candy Incorrect | .83 | .77; .89 | 20.1 | 13; 24 |
| Boredom Money Correct | .91 | .87; .94 | 20.6 | 11; 24 |
| Boredom Money Incorrect | .88 | .83; .92 | 19.9 | 11; 24 |
| Boredom Neutral Correct | .76 | .66; .84 | 20.4 | 12; 24 |
| Boredom Neutral Incorrect | .80 | .71; .87 | 20.1 | 11; 24 |
| Effort Candy Correct | .83 | .77; .88 | 20.1 | 11; 24 |
| Effort Candy Incorrect | .80 | .73; .87 | 19.8 | 10; 24 |
| Effort Money Correct | .90 | .86; .93 | 20.3 | 12; 24 |
| Effort Money Incorrect | .85 | .79; .90 | 19.8 | 12; 24 |
| Effort Neutral Correct | .67 | .54; .77 | 20.1 | 14; 24 |
| Effort Neutral Incorrect | .79 | .71; .85 | 20.2 | 10; 24 |
| Control Candy Correct | .82 | .75; .87 | 20.8 | 12; 24 |
| Control Candy Incorrect | .78 | .70; .85 | 20.7 | 12; 24 |
| Control Money Correct | .84 | .78; .89 | 21.0 | 11; 24 |
| Control Money Incorrect | .85 | .80; .90 | 20.5 | 13; 24 |
| Control Neutral Correct | .81 | .75; .87 | 20.8 | 12; 24 |
| Control Neutral Incorrect | .74 | .63; .82 | 20.6 | 12; 24 |

## Results

Syntax and output for all results, as well as the data, are posted on OSF (https://osf.io/e39as/?view_only=03a768b3ec084a86a4a1ce488ce6d164).

### Preliminary analyses: phenomenology of depletion and boredom conditions

We first examined differences between the effort and boredom conditions in self-report ratings of fatigue, boredom, mood, and effort. A mixed (between: condition X within: block) ANOVA found that for all four variables there was a main linear effect of time, as well as a main effect of condition (see Table 2). There was also a significant linear time by condition interaction for fatigue and mood but not for boredom or effort (see Table 2). Figure 3 illustrates these effects. Participants reported feeling more bored (less interested) as the task went on, and boredom was stronger in the bored condition, supporting the effectiveness of our boredom manipulation. Similarly, as expected, participants in the effort condition exerted more effort throughout the task. These self-report ratings validate our method, confirming that participants exerted greater cognitive effort in the effort condition, and felt more bored in the boredom condition. Participants in the bored condition also reported a sharp decrease in mood throughout the experiment, although participants in the effort condition reported more unpleasant mood. Surprisingly, although participants in the bored condition reported levels of fatigue similar to the effort condition after the first block, they reported more fatigue as time went on. Remaining bored, in other words, might have been phenomenologically more fatiguing than continuously exerting cognitive effort.

**Figure 3.**
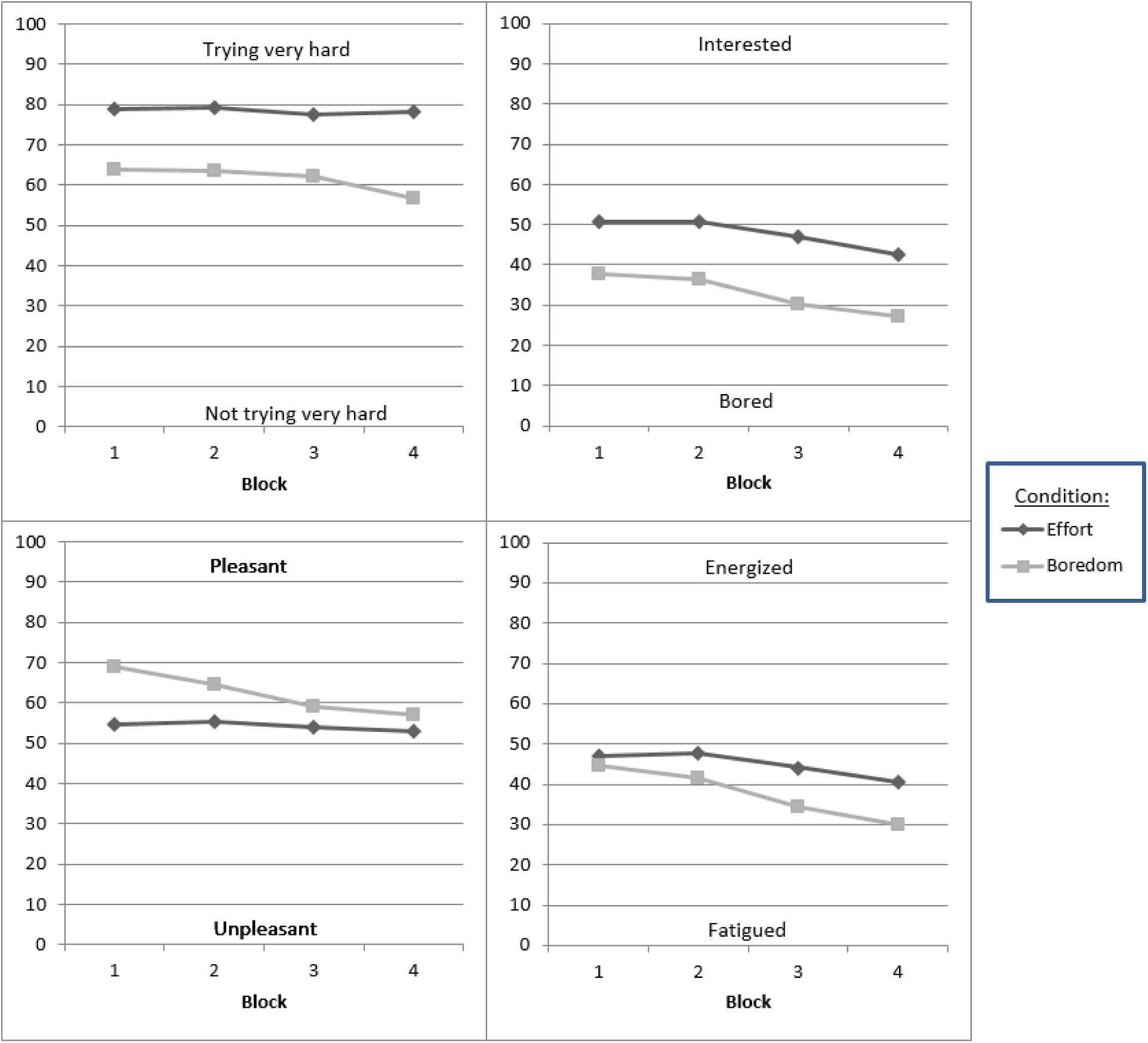
Phenomenology of effort and boredom conditions across blocks. Ratings were made on visual-analog scales ranging from 0-100. The wording for each construct were as follows: Effort (top left) “How hard are you trying to do well on the task” (Not trying very hard – trying very hard); Boredom (top right) “How are you feeling right now?” (Bored-Interested); Mood (bottom left) “What is your mood right now” (Unpleasant-Pleasant); Fatigue (bottom right) “How are you feeling right now” (Fatigued-Energized).

**Table 2.**

| | F | p | $\eta^2$ | 90% CI |
| --- | --- | --- | --- | --- |
| Main effect of time (linear) |  |  |  |  |
| Fatigue | 53.60 | <.001 | .25 | .16; .34 |
| Boredom | 34.29 | <.001 | .18 | .10; .26 |
| Mood | 31.70 | <.001 | .17 | .09; .25 |
| Effort | 8.06 | .005 | .05 | .01; .11 |
| Main effect of condition |  |  |  |  |
| Fatigue | 7.93 | .005 | .05 | .01; .11 |
| Boredom | 23.99 | <.001 | .13 | .06; .21 |
| Mood | 8.97 | .003 | .05 | .01; .12 |
| Effort | 41.20 | <.001 | .21 | .12; .29 |
| Time* condition interaction |  |  |  |  |
| Fatigue | 8.31 | .004 | .05 | .01; .11 |
| Boredom | .77 | .383 | .01 | .00; .04 |
| Mood | 17.65 | <.001 | .10 | .04; .18 |
| Effort | 3.87 | .051 | .02 | .00; .08 |

Note: $\eta^2$ were obtained using the SPSS calculator from Wuensch (2012). 90% CIs are reported (instead of 95%) because $\eta^2$ cannot be negative (see http://daniellakens.blogspot.ca/2014/06/calculating-confidence-intervals-for.html for a clear explanation)

### Main analyses: differences in FN across conditions

We first conducted a between-subject analysis comparing the three conditions on overall FN amplitude. Table 3 reports all the means. There were significant differences among conditions, F(2, 211) = 3.17, p = .04, η_2_= .03, 90%CI[.0004;.0700], although the effect was small in magnitude. To more directly test our hypotheses, we conducted two planned contrasts: one comparing both the effort and boredom conditions to the control condition, and the other comparing the effort and boredom conditions to each other; we expected the former, but not the latter, to be different from one another. The first planned contrast showed that together, the effort and boredom conditions were significantly different from the control condition, *t*(209) = -2.04, *p* = .04, *d* = .30, 95%CI[.01;.59], although again the effect size was only modest in magnitude. A post-hoc examination of the means showed that the effects appeared to be driven by the boredom condition – that is, participants in the boredom condition had the most-negative FN among the three groups (see Table 3). The difference between the boredom and control condition was significant even after correcting the alpha level for the three comparisons (i.e., lower than .05/3, as per a Bonferroni correction), *t*(135) = -2.58, *p* = .011, *d* = .44, 95%CI[.10;.78] ^1^. The second planned contrast, comparing the effort and boredom condition, showed that the two were not different from one another, t(209) = 1.53, p=.13, d = .24, 95% CI [-.09;.57]. Importantly, the direction of the means was such that the boredom condition showed the strongest FN response. Figure 4 illustrates the FN waveforms and topographical maps for each condition.

**Table 3.** Overall FN difference amplitude

|  | <u>N</u> | <u>Mean</u> | <u>SD</u> | <u>95% CI</u> |
| --- | --- | --- | --- | --- |
| Effort | 75 | -.86<sub>ab</sub> | 1.37 | -1.17; -.54 |
| Boredom | 68 | -1.19<sub>a</sub> | 1.37 | -1.52; -.86 |
| Control | 69 | -.64<sub>b</sub> | 1.14 | -.91; .36 |

Note: Values not sharing a subscript are significantly different from each other in post-hoc analyses.

**Figure 4.**
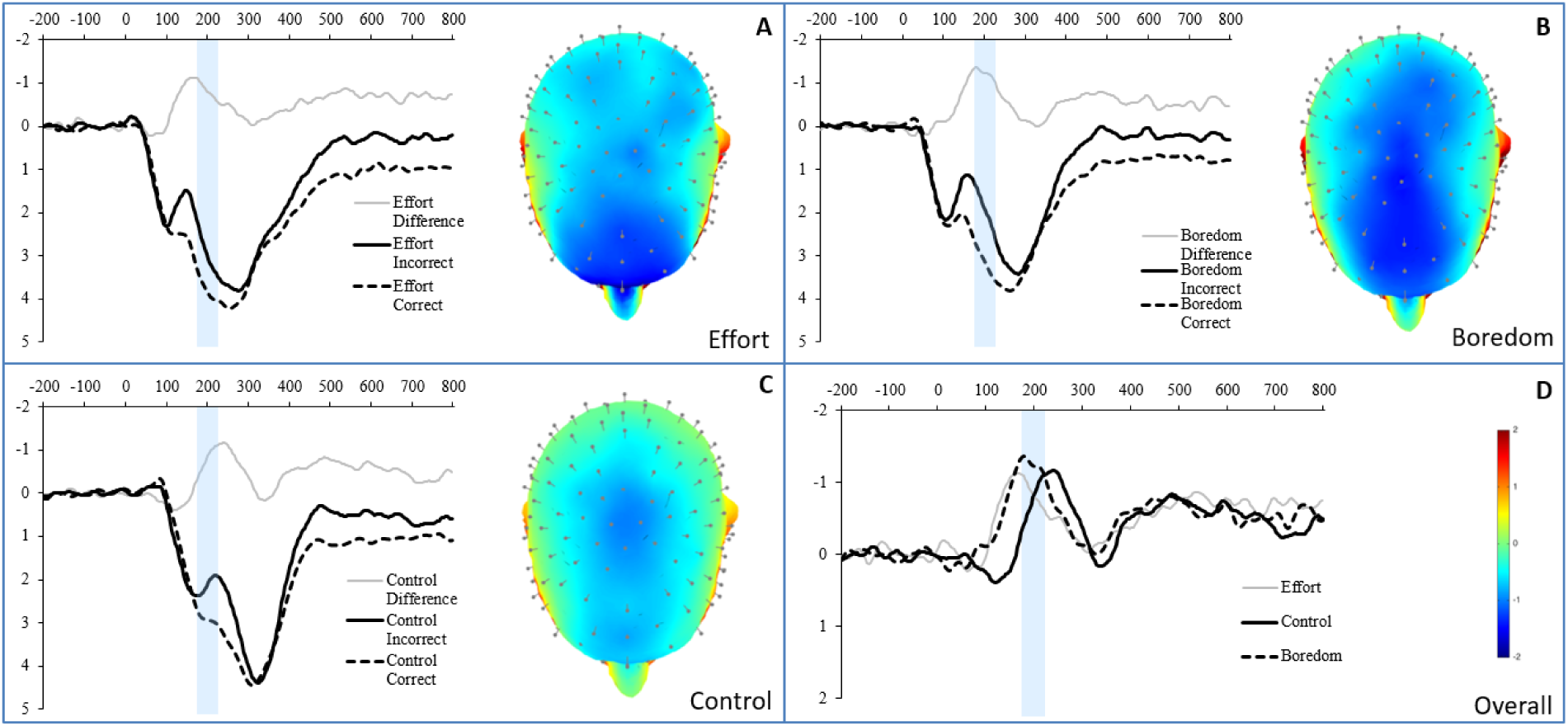
Response-locked ERP waveforms comparing gain (correct) and loss (incorrect) trial waveforms, in the Effort (top left), Boredom (top right), and Control (bottom left) conditions, as well as the waveforms of the difference between gains and loses across the three conditions (bottom right). For each panel, response onset occurred at 0 ms and negative is plotted up. The grey bar illustrates the time interval (175-225ms) used for calculating the value of the FN in the present study. Also shown are topographic maps depicting differences between response to gains and losses in Effort (top left), Boredom (top right), and Control (bottom left) conditions at the 200ms time.

To examine whether the type of reward matters, we ran a 3(between) X 3(within) MIXED model in SPSS. Condition (effort, boredom, control) was entered as a between factor, and type of reward (money, candy, positive feedback) was entered as a within factor. There was a main effect of reward type, F(2,391.60) = 5.88, p = .003, η_2_= .03, 90%CI[.006;.060], and a marginal main effect of condition, F(2,195.89) = 2.94, p = .055, η_2_= .03, 90%CI[.000;.072] but no interaction effect, F(4, 391.58) = .70, p = .593, η_2_= .007, 90%CI[.000;.016]. A post-hoc examination of the means showed that across the types of rewards, the boredom condition (M=- 1.22, 95%CI[-1.53;-.91]) elicited a significantly larger FN that in the control condition (M = - .69, 95%CI[-.99;-.39]), mean difference =.52 95%CI[.10;.96], p = .017). Furthermore, across the three conditions, money elicited a larger FN response (M = -1.19, 95%CI[-1.42;-.95]) than both candy (M = -.90; 95%CI[-1.13;-.67]; mean difference = .29 95%CI[.01;.56], p = .041) and positive feedback (M = -.71, 95%CI[-.95;-.48]; mean difference = .48 95%CI[.20;.75], p = .001); the latter were not significantly different from one another, mean difference = .19 95%CI[- .09;.46], p = .177. However, as there was no interaction, this did not differ across conditions.

### Supplementary analyses: linking phenomenological reports of depletion with FN

To further examine whether self-reports of depletion were related to reward sensitivity, we conducted exploratory analyses testing the correlation between self-reported boredom, fatigue, and effort (averaged across the 4 time points) with reward sensitivity as indexed by the FN. Since these self-report measures were only available for participants in the boredom and effort conditions, only data from these participants (N=143) was used. There was no relationship between these self-report variables and the FN, all *r*s < .1, *p*s> .40.

## Discussion

The present paper examined the effects of cognitive effort and boredom on reward sensitivity, as indexed by the FN. Although in planned contrast analyses we found that participants in both boredom and effort conditions (combined) exhibited a stronger (i.e., more negative) FN response than participants in the control condition, post-hoc analyses and within-subject multilevel analyses showed that this was primarily driven by participants in the boredom condition. That is, the boredom condition led to greater reward sensitivity. Even though previous research has found that people who are generally more prone to experience boredom (i.e., trait boredom) engage in greater reward-seeking and impulsive behaviours, our study was the first to examine the neural consequences of state boredom. Here, we find that boredom, experimentally induced in the laboratory, is associated with heightened brain responses to rewards. This could explain why people who frequently experience boredom act in a more impulsive matter, as sensitivity to reward may lead people to actively seek reward (e.g., Loxton & Dawe, 2001). Interestingly, participants in the boredom condition reported feeling particularly fatigued, more so than those in the effort condition. The phenomenological experiences of boredom need to be further investigated.

Although the boredom condition led to greater reward sensitivity than the control condition, the effort condition was neither different from the boredom nor from the control condition. Contrary to predictions, we did not find that participants who engaged in cognitive effort were more reward sensitive than those who did not exert effort; this despite clear evidence that participants in the effort condition reported exerting effort and were (at least to some extent) mentally fatigued. This differs from past research on reward perception, which has found that mental effort leads to increased approach motivation to rewarding stimuli (Schmeichel et al., 2010) and to higher activation in the orbitofrontal cortex (and greater functional connectivity between the orbitofrontal cortex and the inferior frontal gyrus) in response to food cues (Wagner et al., 2013). It is also at odds with recent experimental research on reward seeking, which has shown that depleted individuals will strive for rewards when these are easy (but not difficult) to obtain (Giacomantonio et al., 2014). Clearly, more research is needed to understand whether effort affects reward processing, and under what conditions this would occur.

Exploratory supplementary analyses showed that neither effort, boredom, mood, nor self-reported fatigue were related to the FN for participants in the effort and boredom conditions. Previous analyses have occasionally linked effort reported on a depleting task with ego-depletion effects observed in the second task in the sequential task paradigm (Dang, 2016), although these effects tend to be quite small. In contrast, other studies have shown a dissociation between perceptions of effort/fatigue and actual fatigue (e.g., Clarkson et al. 2010). This is in line with other research on phenomenology, especially in the area of emotions. Although phenomenal reports of emotion and emotional behaviour are theoretically assumed to cohere, empirical research rarely finds such evidence. While some studies suggest mild coherence, at least in certain situations (e.g., Mauss, Levenson, McCarter, Wilhelm, & Gross, 2005), the vast majority of studies find that emotional response systems do not correlate very highly (e.g., Lang, 1968; Weinstein, Averill, Opton, & Lazarus, 1968). Such lack of coherence, in fact, has led to a broad rethinking of emotion altogether (Barrett, 2006). This is important because it suggests possible dissociations between people’s subjective experience of their affective state and their physiological and behavioral expressions of these states. Future research is thus needed to better understand how phenomenological experiences actually relate to brain and behaviour.

Our results point to the importance of considering the procedures that research on ego depletion uses in the so-called control condition. Indeed, it may be the case that some of the control conditions used in ego depletion studies actually elicit boredom. For example, studies that require participants to read uninteresting texts and cross out letters (e.g., Baumeister et al. 1998), or engage in a repetitive task that does not require mental effort (e.g., Hagger et al. 2016) may all have led to boredom, which we have found to be related with higher subjective fatigue and greater sensitivity to rewards. This may explain the small effects (or lack of effects) found in some depletion studies - the comparison group in some studies may not be a true neutral control. Future studies need to consider the control groups that are used to ensure that boredom is not accidentally induced.

The current study also furthers research on the FN as an index of reward sensitivity (Proudfit, 2015). Using a standard task for eliciting the FN (Proudfit, 2015), we found again that monetary rewards elicit a larger FN response than positive feedback alone (Weinberg et al., 2014). Adding a new condition with another type of tangible, though non-monetary reward (M&M candy), results suggested that an appetitive food-type of reward was not different from the neutral (feedback-only) condition. This may have interesting implications for how rewards are processed in the brain, and especially what is rapidly processed in the brain as rewarding. It is interesting that money, which is a social construct, can elicit greater brain reactivity than food, which should arguably be a more readily salient and immediate reward. More research is needed to understand whether this is indeed the case (i.e., was it jut M&Ms that were not considered rewarding, or food in general), and why and how such a response would have developed.

### Limitations

This study had several limitations. First, despite the large sample size, we might still have been underpowered, if the effect size was smaller than a medium effect size. For example, we only had 50% power to detect a small effect size of f = .15. Alternatively, a more powerful manipulation of effort and boredom may have helped us to see more differentiation between the constructs. Second, we did not have a manipulation check (i.e., measures of fatigue, boredom, etc.) for participants in the control condition; it is thus possible that we had a limited range in the phenomenological measures, which may be why we did not find any relations between phenomenological experiences and reward sensitivity.

Similarly, we do not know how much effort participants actually exerted on the two tasks – it may be that some people disengaged from the effortful task (due to difficulty being too high – see Gendolla, Wright & Richter, 2004), while the boredom task may have led people to exert effort to remain vigilant. This latter possibility is supported by participants’ ratings of their effort, which was still over the midpoint even in the boredom task. Despite the effortful condition being more demanding than the boredom condition, perceptions and motivation may have affected the actual use of effort. Indeed, a rich literature on the motivation intensity theory (Brehm & Self, 1989) describes how multiple processes related to the self (e.g., motivation, ego involvement) can interact with task difficulty to determine the amount of effort that a person will invest in a task (Gendolla & Richter, 2010). Future research is needed to consider these sources of effort and calibrate both tasks, replicating these results with other tasks requiring cognitive effort (and leading to boredom) to ensure generalizability.

A final potential limitation is in the timing of our FN component. Typically the FN is seen between 250 and 350ms. Our earlier FN is somewhat unusual. However, we used a more objective approach (collapsed localizer) than simple visual inspection of the waveforms to determine our window, the waveform morphology is consistent with studies of the FN, and the component obtained highly dependable estimates with as few as 10 trials. Thus, while early for a typical FN, we feel are measuring the correct ERP component.

## Conclusion

The present study was the first to examine neural sensitivity to rewards following cognitive effort in the general population. Importantly, we contrasted cognitive effort to boredom, an affective state that could be expected to increase sensitivity to rewards. Results suggest that while bored participants were indeed more responsive to rewards than participants in a control condition, this was not the case for participants who had exerted cognitive effort. That is, we found little support for our hypothesis that cognitive effort would increase reward sensitivity. Interestingly, exploratory analyses also found that participants in the boredom condition reported greater subjective fatigue than participants in the cognitive effort condition, suggesting that boredom can be fatiguing. Together, these results shed new light on the phenomenological and cognitive consequences of experiencing boredom.

## Acknowledgments

This research was supported by funding from the Brigham Young University College of Family, Home, and Social Sciences to M. Larson. M. Inzlicht was funded by the Social Sciences and Humanities Research Council of Canada (SSHRC; grant #435-2014-0556).

## Footnotes

1 d and confidence interval is calculated based on t-test directly comparing the two conditions.

